# CCAR-1 has a novel role in regulating the *Caenorhabditis elegans* germline

**DOI:** 10.1101/2021.07.06.451293

**Authors:** Doreen I. Lugano, Andrew Deonarine, Margaret A. Park, Sandy D. Westerheide

## Abstract

The Cell Division Cycle and Apoptosis Regulator (CCAR) protein family members are putative transcription regulators that have been characterized for modulating the cell cycle, apoptosis, metabolism, and the heat shock response. Mammals have two CCAR family members, CCAR1 and CCAR2/DBC1, that evolved from the founding family member CCAR-1 that is expressed in *Caenorhabditis elegans*. Mammalian CCAR2, the most well-studied family member, has been shown to regulate genes involved in metabolism in cultured cells. However, the regulation of gene expression by CCAR family members at an organismal level is unknown. Here, we use whole transcriptome RNA sequencing to examine the effects of CCAR-1 on gene expression in *Caenorhabditis elegans.* We show that CCAR-1 regulates germline transcription, reproduction, lifespan, and DNA-damage induced apoptosis. This study shows the role of CCAR-1 in vital physiological functions in the *C. elegans* germline that have not been investigated before.

## 2. INTRODUCTION

The Cell Division Cycle and Apoptosis Regulator (CCAR) proteins CCAR1 and CCAR2 regulate a wide variety of physiological processes (Brunquell et al. 2014a). Mammalian CCAR1 and CCAR2 are predicted through phylogenetic analysis to have evolved from the *Caenorhabditis elegans* CCAR-1 (Brunquell et al. 2014b). Like mammalian CCAR proteins, *C. elegans* CCAR-1 has been shown to regulate a diverse array of functions, including epigenetic modifications, stress responses, and RNA splicing (Brunquell et al. 2018b; Brunquell et al. 2014b; Fu et al. 2018).

Mammalian and *C. elegans* CCAR proteins have 30% sequence similarity and share similar domains (Brunquell et al. 2014b). CCAR family domains of note include an S1-like putative RNA binding domain, a nuclear localization sequence, a leucine zipper domain that may allow for protein-protein and protein-DNA interaction, a Nudix domain that is predicted to bind to metabolites, an EF-hand domain, and coiled-coil domains that may allow for protein-protein interaction. Additionally, mammalian CCAR2 and *C. elegans* CCAR-1 both have a SAP domain, which is predicted to allow for DNA binding.

These functional domains, along with the fact that these proteins have been localized to the nucleus, support the hypothesis that CCAR proteins may regulate gene expression both directly and indirectly. It is possible that CCAR proteins bind to DNA directly and function as transcription factors. CCAR proteins may also affect transcription through the modulation of transcription factors. For example, human CCAR2 directly interacts with steroid hormone receptors, affecting their transcriptional functions (Fu et al. 2009). CCAR proteins are also known to regulate transcription epigenetically through effects on the deacetylase SIRT1 (Kim et al. 2008; Zhao et al. 2008).

Transcriptome-wide RNA sequencing studies in mammalian tissue culture studies have shown that CCAR family members regulate gene expression of genes involved in metabolism. Studies using RNAi against CCAR1 or CCAR2 in A549 human lung cancer cells showed that this protein affected the expression of steroid hormone receptors, many of which are involved in cell metabolism. This makes sense, as CCAR proteins interact with steroid hormone receptors (Wu et al. 2014). Additional studies with CCAR2 RNAi performed in HEK 293 human embryonic kidney cells confirmed the involvement of CCAR2 in regulating genes involved in metabolism (Basu et al. 2020; Close et al. 2012).

Are metabolic genes the only gene category affected by CCAR proteins? To answer this question, we employed *C. elegans* as a whole animal model organism. We chose the worm as a model for our studies as there is only one CCAR family member, a CCAR-1 deletion strain is available, and we can assess gene expression in multiple tissues. Through RNA-sequencing, we demonstrate that the deletion of CCAR-1 affects the regulation of many gene categories. As expected, a top category of regulated genes are metabolism genes from the insulin signaling-related pathways. However, using our *C. elegans* model, we have uncovered an additional category CCAR-1-regulated genes: germline genes. Interestingly, we find that CCAR-1 RNAi increases stress-induced apoptosis in the germline and leads to decreased progeny numbers. Additionally, we find that CCAR-1 RNAi increases lifespan, an effect that is dependent on the ability of CCAR-1 to regulate the germline. Thus, we have identified germline genes as a novel category of genes regulated by CCAR-1, providing a mechanism by which CCAR-1 can regulate lifespan in *C. elegans*.

## 3. METHODS

### 3.10 *C. elegans* strains and maintenance

The following strains were used in the study: Wild-type Bristol N2, CCAR-1Δ VC1029 (gk433), CCAR-1Δ MT7019 (n2667), CCAR-1Δ DM1153 (ra14), CCAR-1Δ DM1154 (ra5), PRG-1Δ WM161 (tm872), ACT-5∷YFP strain WS2170 (unc-119(ed-3)), GLP-1Δ CF1903 (e2144), and SDW080 (peft∷CCAR-1∷GFP∷3xflag). The CCAR-1Δ VC1029 (gk433) mutant strain was outcrossed three times to our N2 wild-type strain to generate the SDW040 strain. N2, VC1029, MT7019, DM1153, DM1154, WM161, WS2170, and SDW080 strains were maintained at 20 °C on standard NGM plates seeded with *Escherichia coli* OP50-1. Synchronous populations of nematodes were obtained by bleach synchronization and plating for 19 hrs on NGM plates without food. GLP-1Δ CF1903 (e2144) was maintained at 16°C until bleach synchronization where the L1s were then split and grown at either 16°C or 25°C.

### 3.11 RNA preparation for RNA sequencing

RNA samples were prepared using Trizol reagent (Ambion. The samples were sent to the Brigham Young University DNA sequencing center. RNA libraries were prepared using the TruSeq Total RNA kit (Illumina, Inc.) with a Ribo-zero gold kit (Illumina, Inc.) to remove rRNA and mitochondrial RNA transcripts.

### 3.12 RNA-sequencing analysis

HISAT and STRING-Tie were used in the alignment and assembly of raw FASTQ files from the RNA sequencing. The R program, Ballgown package, was used to calculate reads per kilobase of the transcript, per million mapped reads. The same package was used to calculate the statistics of the data set, which include the p-value, the q value which has Benjamini-Hochberg correction for multiple testing, and the log2 fold change of genes.

### 3.13 Heat map generation

The heatmaps were generated using R Studio. The Bioconductor package with the gplot library, and the heatmap.2 function was utilized.

### 3.14 Venn Diagram analysis

Bioinformatics and genomics website http://bioinformatics.psb.ugent.be/cgi-bin/liste/Venn/calculate_venn.htpl and LucidChart were used to construct Venn diagrams. Venn diagrams were generated for significantly altered mRNA for each condition tested (p-value <0.05).

### 3.15 Brooding Assay

The wild-type Bristol N2, CCAR-1Δ VC1029 (gk433), CCAR-1Δ MT7019 (n2667), CCAR-1Δ DM1153 (ra14), CCAR-1Δ DM1154 (ra5), were used in the brooding assays. Synchronized animals were grown on OP50 plates until the L4 molting stage in both 20°C. Individual worms were transferred to 12 well plates to assess daily brooding amounts. The worms were moved to a new well each day, and the number of progeny was recorded. The experiments were repeated in triplicate with n=6 for each trial. The same procedure was repeated for all the strains in EV and sir-2.1 RNAi plates.

### 3.16 qRT-PCR

qPCR was performed to determine knockdown of sir-2.1 by RNAi in N2 and CCAR-1 Δ. Synchronized animals were grown on control (EV) and sir-2.1 RNAi plates until L4 molting stage at 20°C. Worms were collected, and RNA extraction was done using Trizol reagent (Ambion) followed by cDNA synthesis using the SuperScript III reverse transcriptase kit (Invitrogen) according to the manufacturer’s instructions. cDNA was diluted to 50ng/μl and used as a template for qRT-PCR. qRT-PCR and the normalization was done as previously described (Brunquell et al. 2017).

### 3.18 Fluorescence imaging

WS2170 (act-5∷YFP strain) was synchronized and fed with control (EV), CEP-1, CED-4, and CCAR-1 RNAi from the L1 to L4 developmental stage. Worms were irradiated at L4 with 400uJ/m3 UV using a Stratalinker. Treated worms were imaged for the presence/absence of act-5∷YFP ‘halos’ 24 hours post-radiation using a Keyence fluorescence microscope.

SDW080 animals were anesthetized with 10mM levamisole and photographed using a Keyence fluorescence microscope.

### 3.19 Lifespan Assays

Lifespan assays were conducted on wildtype N2 and GLP-1Δ CF1903 (e2144) at both 16°C and 25°C with forty worms per condition in biological triplicates. Animals were transferred to fresh plates daily for five days to avoid progeny contamination. Adult worms were scored every other day and counted as dead upon no response to poking with a platinum wire. Survivability was plotted using Graphpad Prism v.6 (Graphpad software www.graphpad.com) and statistical analysis done by log rank (Mantel-Cox) test.

### 3.20 Data Availability

*Gene expression data are available* in the SRA database (Accession Number SUB8460771).

## 4. RESULTS

### 4.1 Germline genes are regulated by CCAR-1

We used whole transcriptome RNA-sequencing to examine gene regulation by *C. elegans* CCAR-1. The CCAR-1 deletion mutant strain VC1029, which contains a deletion of the first three exons of the *ccar-1* gene resulting in a null mutant (Fu et al. 2018), was outcrossed three times with our laboratory wildtype N2 strain to create strain SDW040. RNA samples were then generated and sequenced from larval stage four (L4) wildtype (N2) and CCAR-1Δ (SDW040) worms in biological duplicates. The top two gene categories regulated by *ccar-1* deletion were found to be germline genes and DAF-2/SIR-2.1-regulated genes, at 24.1% and 20.7%, respectively (Fig. 1A). The DAF-2/SIR-2.1-regulated gene category was expected, as previous work in both *C. elegans* and mammalian cells has linked CCAR family members with SIR-2.1 and SIRT1, respectively (Kim et al. 2008; Zhao et al. 2008; Raynes et al. 2013; Brunquell et al. 2018b). However, a role for CCAR-1 in the regulation of germline genes has not been previously reported. The germline-specific gene categories from our dataset included piRNA/21U-RNA genes, P-granule-associated protein genes, and male-spermatogenic genes (Fig. 1B).

**Figure 1:**
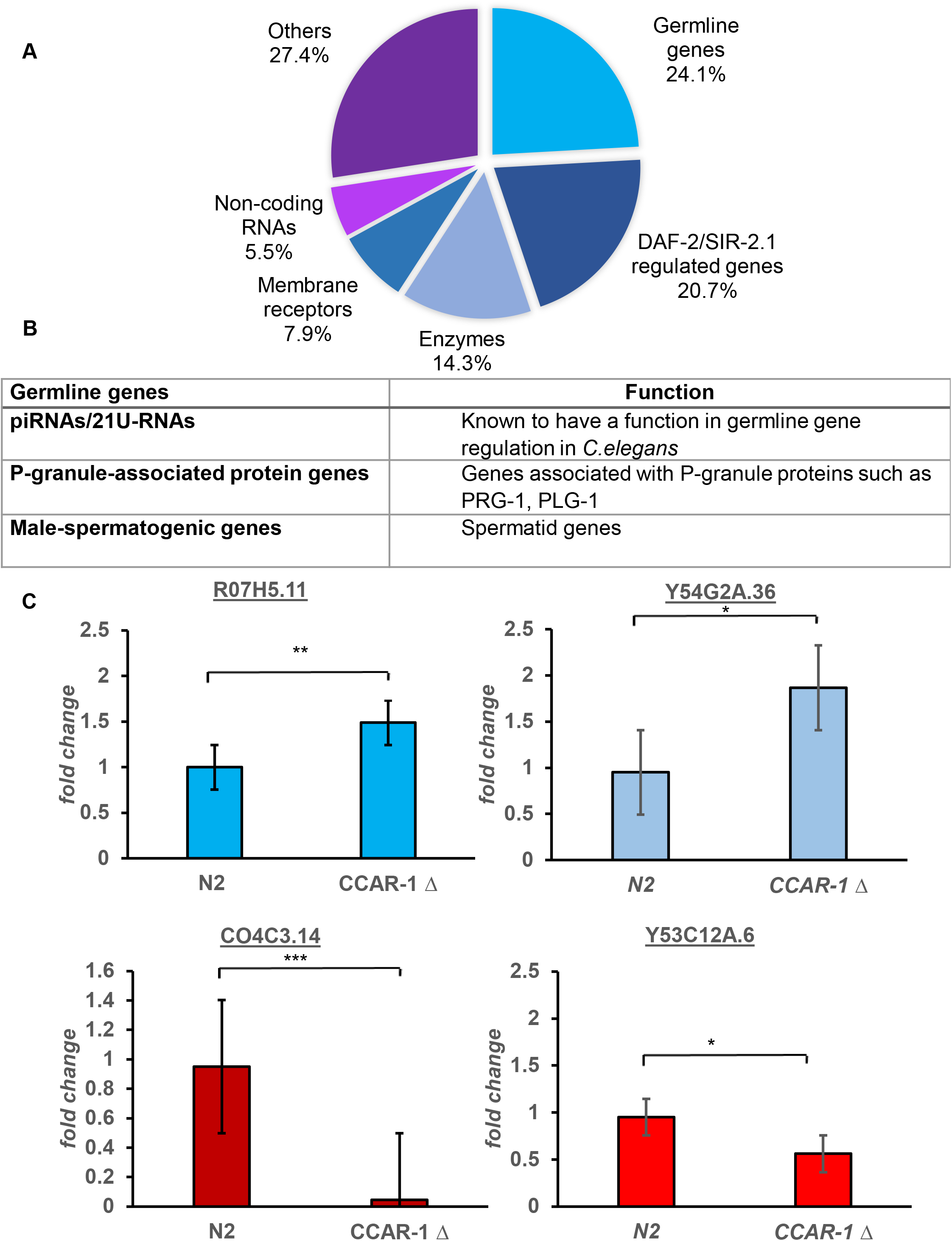
Germline genes are enriched in the gene set regulated by CCAR-1. A. Pie chart showing the genome-wide categories of genes transcriptionally regulated by CCAR-1. RNA sequencing was performed using total RNA samples from N2 (control) and CCAR-1Δ (SDW040) strains. The largest category of regulated genes from the RNASeq analysis are germline genes (24.1%) followed DAF-2/SIR-2.1-regulated genes (20.7%). B. The subcategories of germline-related genes found in our RNASeq dataset are listed. C. qRT-PCR validation of the RNASeq data for genes that are upregulated (blue) or downregulated (red) in the CCAR-1Δ (SDW040) strain as compared to control N2 worms. Significance was determined using the Student’s T test, where * p<0.05, ** p < 0.01, *** p < 0.001

To validate our findings, we then used qRT-PCR on a couple of the top-most downregulated and upregulated germline genes (Fig. 1C). We tested the genes R07H5.11 and Y54G2A.36 from the upregulated gene dataset and F14H8.8 and C04C314 from the downregulated gene dataset and found that the qRT-PCR results were consistent with the RNA-sequencing results. We thus conclude that CCAR-1 regulates the expression of germline-enriched genes, amongst other genes, in *C. elegans.*

### 4.2 CCAR-1 is localized in the germline

As CCAR-1 regulates germline gene expression, we hypothesized that the CCAR-1 protein might be highly expressed in the germline. The CCAR-1 protein has been shown to localize throughout the various cell types of the worm (Fu et al. 2018), but its localization in the germline has not been previously documented. To test for CCAR-1 protein localization, we constructed a peft∷CCAR-1∷GFP∷3XFLAG strain (SDW082) and found that CCAR-1 is expressed in the germline (Fig. 2). CCAR-1 is expressed in the vulva, the oocytes and the mitotic/late pachytene region, supporting its role in germline gene regulation.

**Figure 2:**
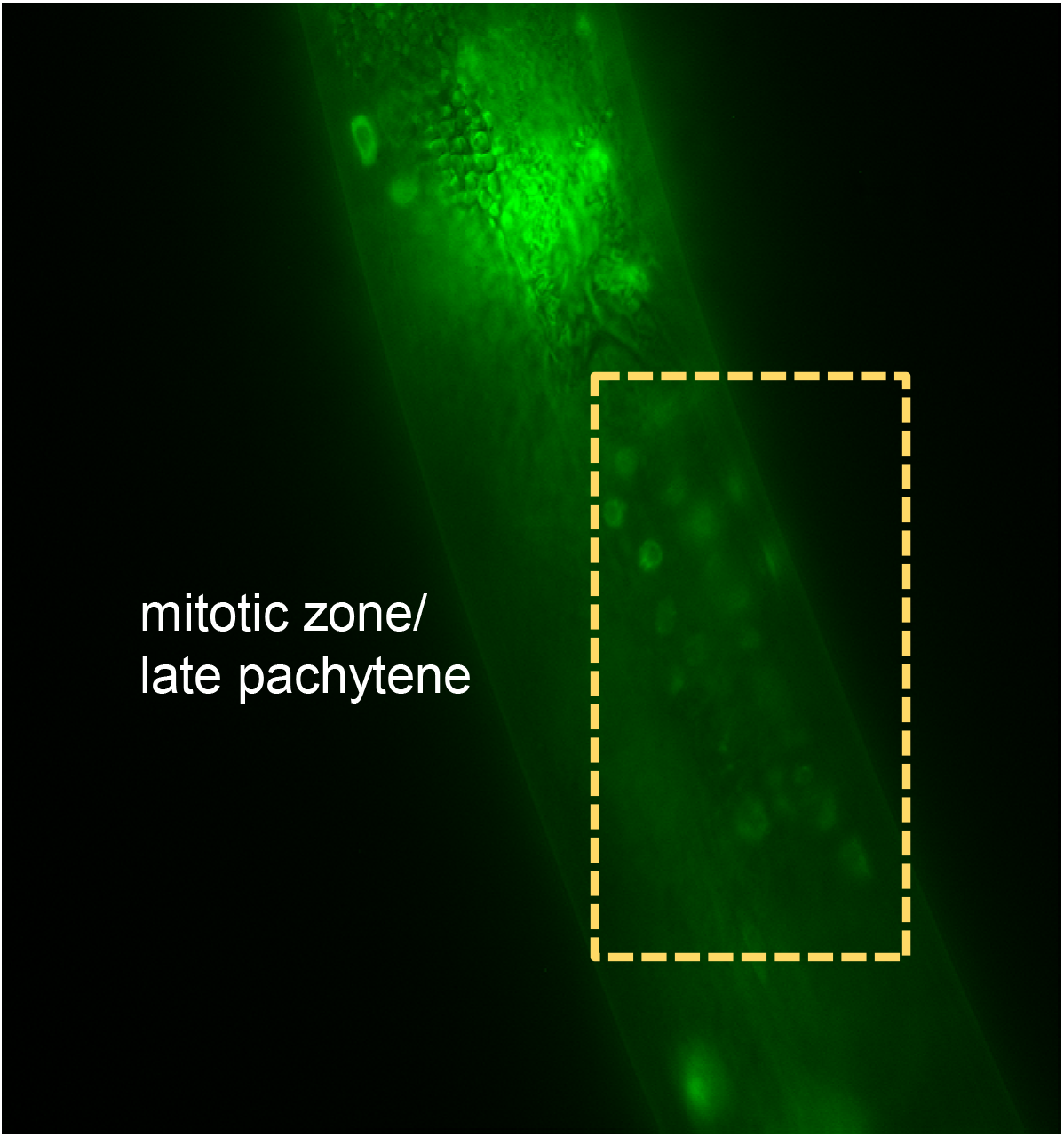
CCAR-1 is localized in the germline. peft∷CCAR-1∷GFP∷3XFLAG animals were anesthetized and photographed using fluorescence microscopy. CCAR-1 is expressed in various germline cells including the vulva, the oocytes and the cells of the mitotic zone/pachytene region.

### 4.3 CCAR-1 protects germline cells from DNA damage-induced apoptosis

Mammalian CCAR1 has been implicated in regulating apoptosis (Francois et al. 2012; Kanno et al. 2019; Muthu et al. 2015; Ou et al. 2014). In *C. elegans*, apoptosis occurs during development and in the gonad of adult hermaphrodites. As we found that CCAR-1 is important in regulating germline genes, we wondered whether it may also be important in regulating germline apoptosis.

During apoptosis, cell corpse engulfment is carried out by neighboring cells, which involves substantial rearrangement of the actin cytoskeleton as it surrounds the dying cells (Povea-Cabello et al. 2017). We thus used a transgenic strain expressing the actin isoform ACT-5 fused to YFP (ACT-5∷YFP) in the sheath cells to highlight apoptotic cells (Lant and Derry 2013; Kinchen et al. 2005). We compared worms treated with empty vector (EV) RNAi as compared to CCAR-1 RNAi to visualize any differences in actin “halos” that appear around early apoptotic cells after treatment with UV damage.

Transgenic ACT-5∷YFP (WS2170) worms were bleach synchronized and placed on RNAi plates until development into L4. L4 worms were then UV irradiated (400uJ/m3) using a Stratalinker and assessed for DNA damage-induced apoptosis 24 hrs later. We found that in the empty vector control RNAi-treated worms (EV) there was an occurrence of several ACT-5∷YFP “halos” signifying the expected UV damage-induced apoptosis (Fig. 3A). As expected, knockdown of either CEP-1/p53 or CED-4/Apaf-1, two key regulators of apoptosis (Lant and Derry 2013; Kinchen et al. 2005), inhibited the production of the UV-induced apoptotic “halos”. Interestingly, upon knockdown of CCAR-1, we see an increase in UV-induced ACT-5∷YFP “halo”’ as compared to control, signifying that CCAR-1 functions in protecting against UV damage-induced apoptosis (Fig. 3A). Data quantification indicates an approximate three-fold increase in actin “halos” with CCAR-1 RNAi as compared to control RNAi (EV) (Fig. 3B). Altogether, these data demonstrate a novel role for CCAR-1 in germline apoptosis.

**Figure 3:**
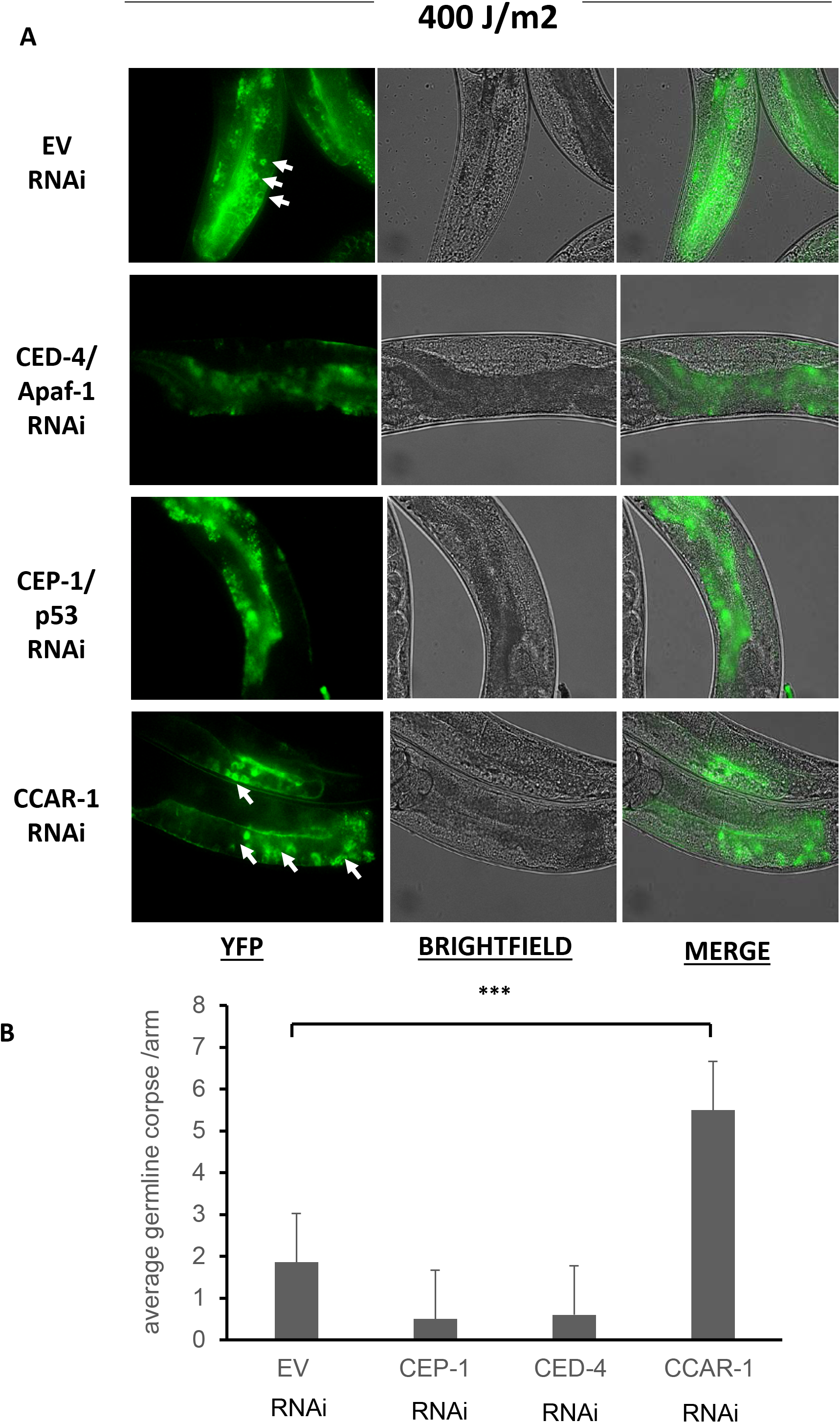
CCAR-1 protects germline cells from UV damage-induced apoptosis. A. ACT-5∷YFP worms expressing actin fused to YFP in the gonad sheath cells were treated with control (EV), CED-4/Apaf-1, CEP-1/p53, and CCAR-1 RNAi prior to exposure to a UV dose of 400J/m2. Images were taken 24 hrs after the exposure. YFP, brightfield and merged images are used to visualize germ cell corpses, which are represented as actin “halos” during early-stage apoptosis and indicated by white arrows. B. Quantification of the number of apoptotic actin “halos” from Figure 3A. There is significant change (p<0.001) between EV and CCAR-1 apoptotic ‘halos’.

### 4.4 CCAR-1 is required for normal levels of progeny production

To further investigate a functional role for CCAR-1 in the germline, we assessed progeny production. We compared wildtype N2 worms with our CCAR-1Δ strain (SDW040) in a brooding assay and counted the number of progeny until day three of adulthood (Fig. 4). To validate our results, we used three additional CCAR-1 mutant strains available from the CGC: MT7019, DM1153, and DM1154. Our results show that CCAR-1 is required for normal levels of progeny production, as progeny levels in the mutants dropped by 2.5-4-fold in the various CCAR-1Δ strains.

**Figure 4:**
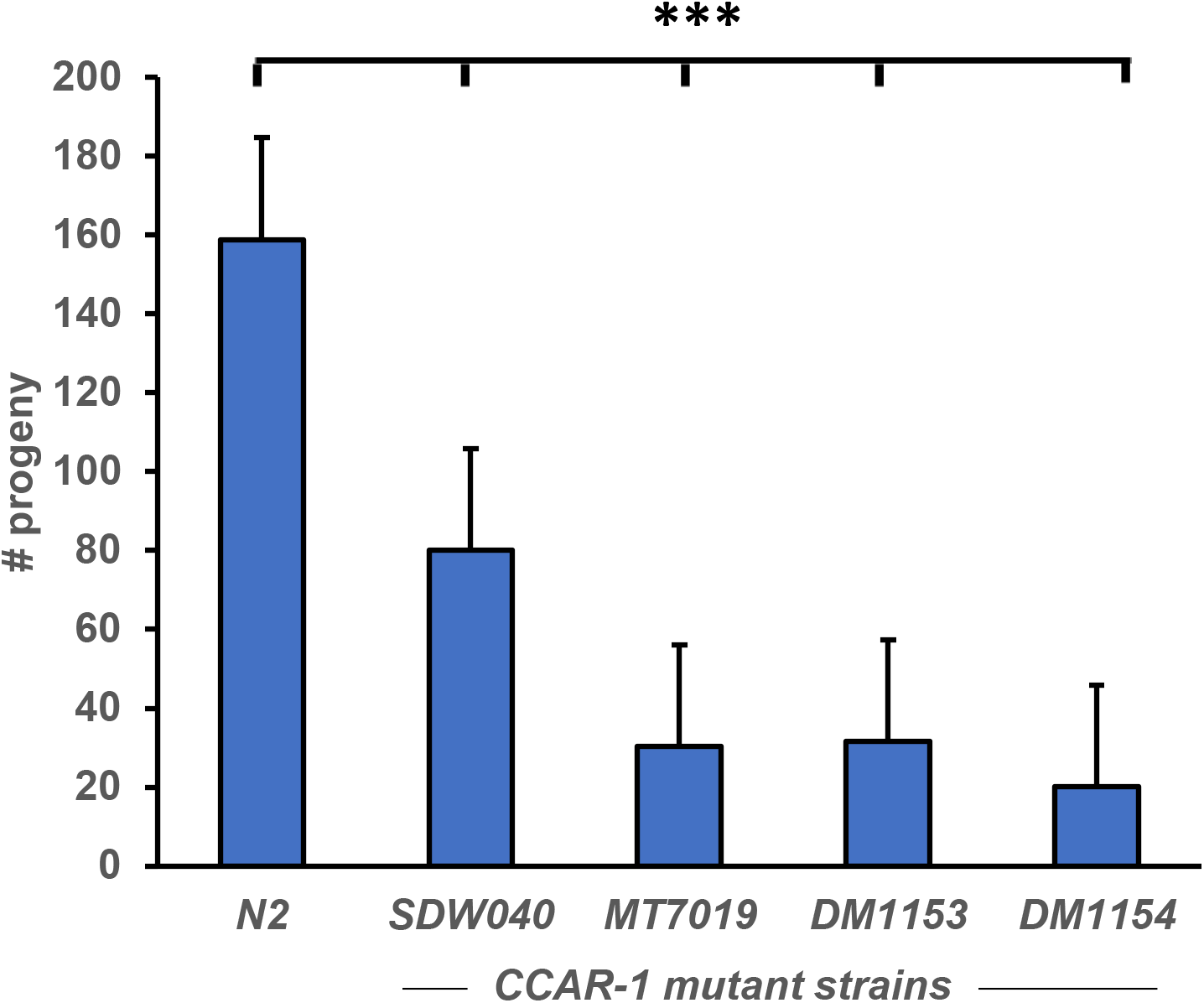
CCAR-1 is required for normal levels of progeny production. The graph shows a brooding assay in which the N2 control strain is compared to various CCAR-1 mutant allele strains (SDW040, MT7019, DM1153 and DM1154) to determine the effects of CCAR-1 on the number of progeny.

### 4.5 CCAR-1 regulates longevity in a manner that is dependent on the presence of a germline

In a previous study, we showed that CCAR-1 RNAi increased *C. elegans* longevity (Brunquell et al. 2018a). The germline in *C. elegans* is known to affect longevity (Brunquell et al. 2018b; Greer et al. 2010; Lin et al. 2001). Therefore, to test whether the effect of CCAR-1 on longevity is dependent on the ability of CCAR-1 to regulate progeny number, we performed lifespan assays using a temperature-sensitive strain in which we could turn off progeny production. GLP-1Δ (CF1903 (e2144)) worms display a temperature-sensitive loss of the germline at 25°C, but not at the permissive temperature of 16°C. We compared the lifespans of wildtype (N2) versus GLP-1Δ (CF1903 (e2144)) worms treated with and without CCAR-1 RNAi (Fig. 5). The worms were scored every other day starting at day 1 of adulthood for survival, and dead worms were scored when non-responsive to poking with a platinum wire. At the permissive temperature of 16°C, both N2 and GLP-1Δ worms showed no change in survival between the control and CCAR-1 RNAi groups, with a mean survival of thirteen days in all cases (Fig. 5A). Thus, CCAR-1 does not affect lifespan at 16°C in the presence of the germline. At 25 °C, wildtype N2 worms fed control RNAi had a mean survival of three days, whereas N2 worms fed CCAR-1 RNAi had a mean survival of five to seven days (Fig. 5B). As expected, based on our previous work (Brunquell et al. 2018a), CCAR-1 RNAi affects lifespan at 25 °C. However, this effect on lifespan extension was lost in the GLP-1Δ worms grown at this temperature (Fig. 5B). At the restrictive temperature of 25 °C in which the germline is absent, GLP-1Δ worms fed control RNAi had a mean survival of eight days, whereas CCAR-1 RNAi fed worms had a mean survival of six days (Fig. 5B). Therefore, these data suggest that decreased CCAR-1 expression enhances longevity through decreasing progeny production (Fig 6).

**Figure 5:**
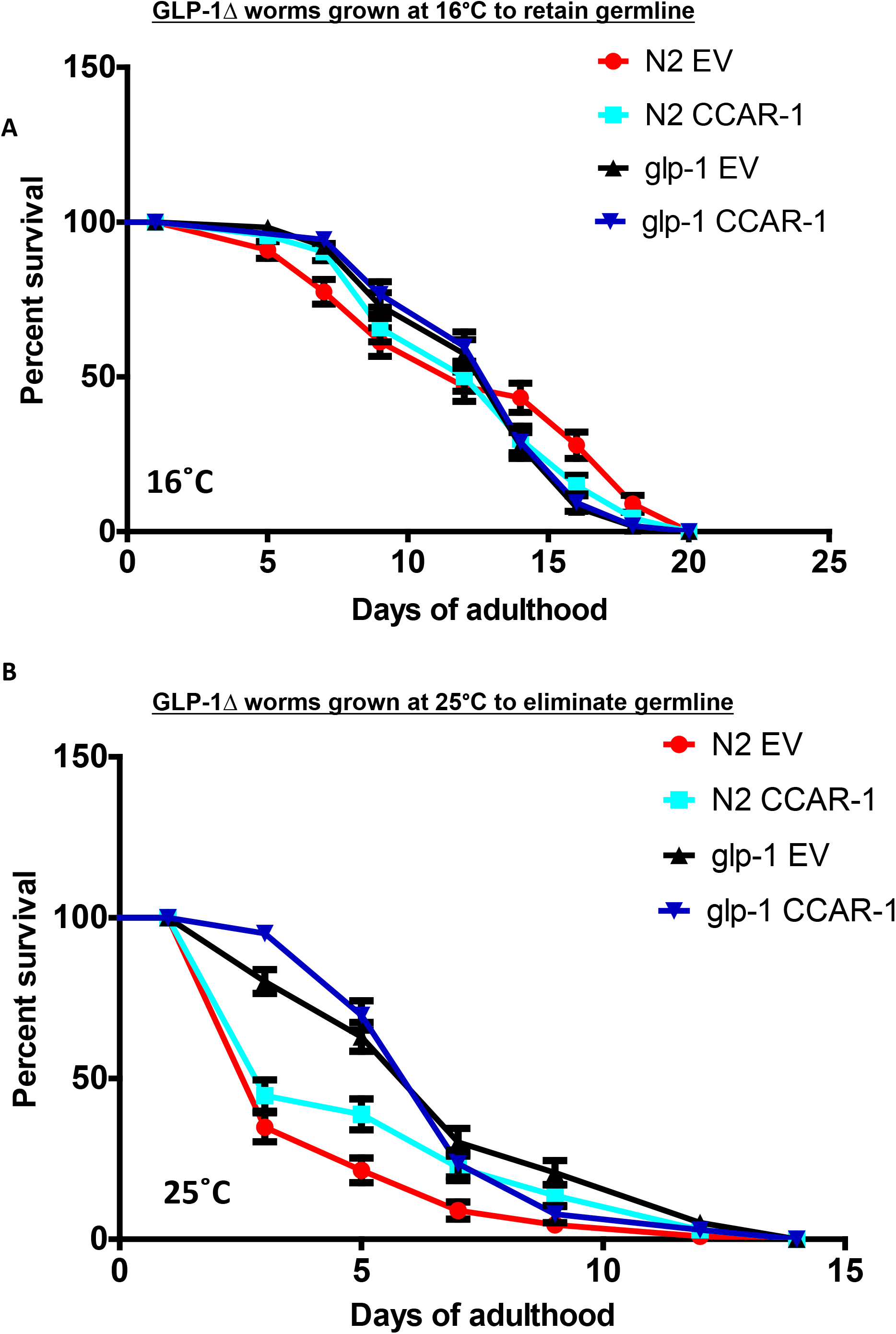
CCAR-1 regulates longevity in a manner that is dependent on the presence of a germline. A. Lifespan analysis was performed at 16°C in wild-type (N2) worms and in GLP-1Δ worms fed control RNAi or ccar-1 RNAi. GLP-1Δ worms grown at this permissive temperature contain a germline. B. Lifespan analysis was performed at 25°C in wild-type (N2) and GLP-1Δ worms fed control RNAi or ccar-1 RNAi throughout lifespan. GLP-1Δ worms grown at this restrictive temperature lack a germline. For (A-B), worms were scored every other day for survival, and significance was determined using the Log-rank (Mantel-Cox) Test.

**Figure 6:**
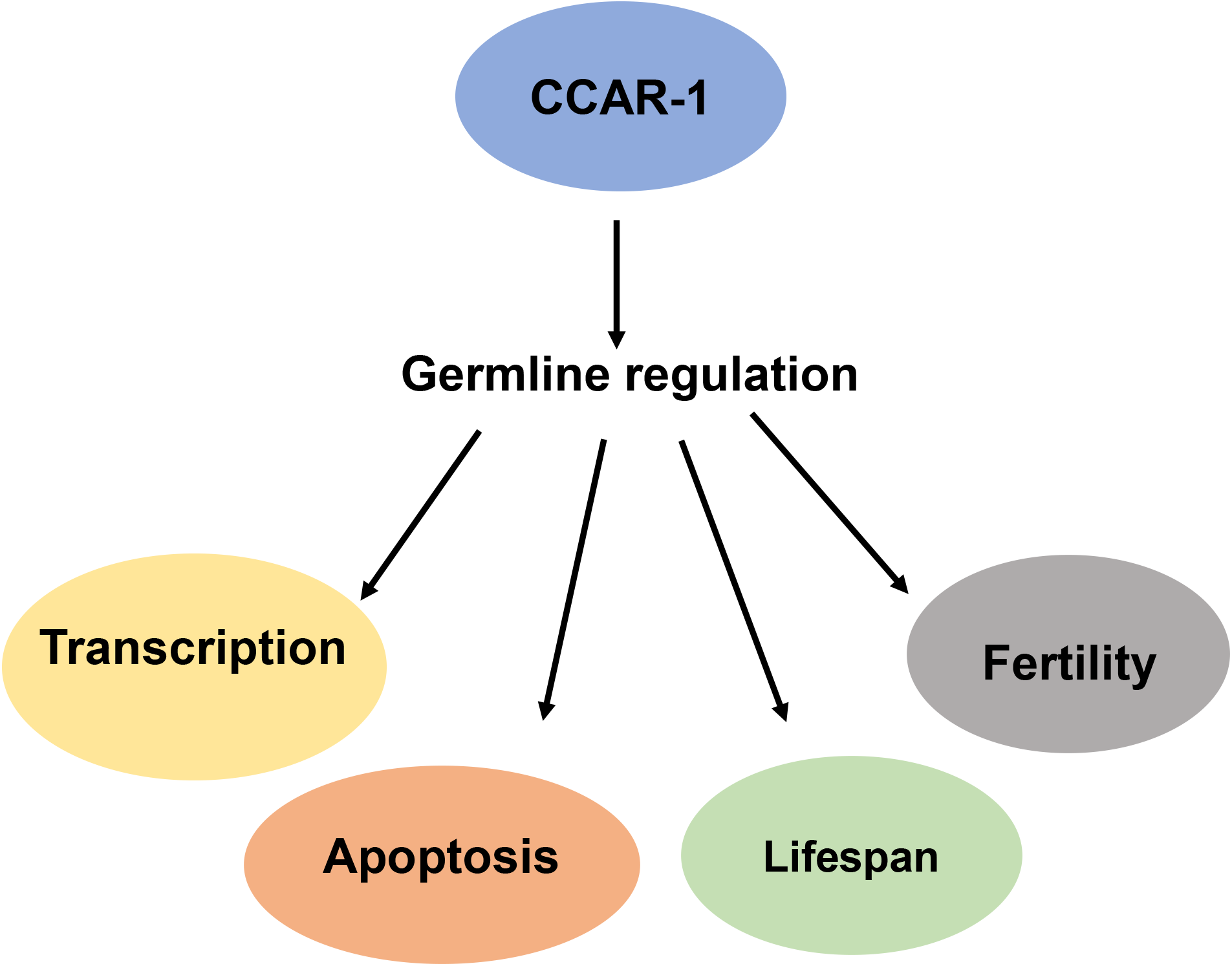
Proposed model for CCAR-1 functions in the germline. We show in this work that CCAR-1 regulates germline gene expression and protects from UV-induced apoptosis in the germline. Reductions in CCAR-1 expression lead to decreased progeny numbers and an enhanced lifespan.

## 5. DISCUSSION

Here, we reveal the genome-wide effect of CCAR-1 on gene expression in *C. elegans.* Previous mammalian tissue culture studies have linked the CCAR family of proteins to the regulation of genes involved in metabolism. We used a strain in which the single worm CCAR family member, CCAR-1, is deleted in these studies. We found that in addition to having a prominent role in regulating metabolic genes, CCAR-1 also controls germline-specific gene expression. CCAR-1 expression is required to protect germline cells in the worm against UV-induced apoptosis and is required for normal levels of progeny. Additionally, CCAR-1 regulates longevity in a manner that is dependent on the presence of a germline.

Our RNA-seq transcriptional dataset shows that the two major categories of genes regulated by the deletion of CCAR-1 in the worm are DAF-2/SIR-2.1-regulated genes, and germline-linked genes. The DAF-2/SIR-2.1-regulated gene category was expected, as human CCAR2 binds to the catalytic domain of SIRT1, forming a stable complex that inhibits SIRT1 deacetylase activity (Kim et al. 2008; Zhao et al. 2008). SIR-2.1 modulates the well-studied DAF-2/DAF-16 signaling cascade (Wang and Tissenbaum 2006; Berdichevsky et al. 2006). Additionally, our previous studies show that CCAR-1 affects the regulation of the heat shock response in a SIR-2.1-dependent manner (Raynes et al. 2013; Brunquell et al. 2018a).

The regulation of germline-related genes by CCAR-1 is a novel finding. The germline-specific gene categories from our dataset included piRNA/21U-RNA genes, P granule-associated protein genes, and male-spermatogenic genes. In the germline of *C. elegans* and other species, piRNAs work together with PIWI argonaute proteins to repress transposable elements from causing detrimental effects in the next generation (Bagijn et al. 2012). In *C. elegans*, piRNAs have also been shown to have a function in spermatogenesis, maintaining germline cells and cell totipotency (Wang and Reinke 2008; Weick and Miska 2014). Future work can help to determine whether CCAR-1 regulates the transcription of these piRNA genes, and/or if it is involved in regulating piRNA processing.

P granules are protein- and RNA-containing membrane-less organelles which are important in *C. elegans* germline development. Interestingly, P granule-associated protein-coding transcripts are another category of genes regulated by CCAR-1.P granule assembly depends on self-interaction domains that are present in the PGL P granule scaffolding proteins. P granules are found throughout the life cycle of worm germline cells, and proteins associated with P granules have a critical role in germline cell differentiation during post-embryonic development (Seydoux 2018). As P granules sit on the cytoplasmic side of nuclear pores, most mRNAs transcribed in germ cells pass through a P granule on their way to the cytoplasm, consistent with a role of these granules in mRNA surveillance. Our finding that CCAR-1 regulates the expression of both piRNAs, and P granule-associated proteins suggests that CCAR-1 may regulate mRNA surveillance in the *C. elegans* germline.

The last category of germline-related genes identified to be regulated by CCAR-1 are male-spermatogenic genes. Previous studies have reported that there are precise sex-specific expression levels of spermatogenic and oogenic functional genes in the male and hermaphrodite germline cells. (Ebbing et al. 2018) With this distinction, several male-spermatogenic genes were shown to be regulated by the deletion of CCAR-1 including spermatid genes. Thus, by using a whole animal model, we have identified a novel role for CCAR-1 in regulating germline genes.

Given that CCAR-1 regulates genes that are important in the germline, our results that extend CCAR-1 protein expression to the germline make sense. While our work is the first to identify CCAR-1 protein expression in the germline, previous high-throughput experiments have shown that the CCAR-1 mRNA is present in the *C. elegans* germline. From a Serial Analysis Gene Expression (SAGE) high-throughput analysis of the hermaphrodite germline, *ccar-1* appeared to be one of 1063 genes enriched in the germline as compared to the soma (Wang et al. 2009). This gene was also identified as differentially expressed between wildtype N2 and GLP-4 worms, which are lacking a fully functional germline (Reinke et al. 2004). Therefore, both *ccar-1* mRNA and CCAR-1 protein are localized to the germline, allowing the possibility for a role in regulating germline gene expression.

Interestingly, P granules and their resident PGL proteins are linked to germline apoptosis. In the *C. elegans* germline, and in the germlines of other organisms, several hundreds of cells die by apoptosis due to physiological signals, genotoxic stress and/or bacterial infection (Gartner et al. 2008). Here, we show that knockdown of CCAR-1 not only regulates germline-specific genes, but also increases the formation of apoptotic germline cells following DNA damage induction. Previous work has shown that PGL-depleted germ cells are selectively committed to apoptosis (Min et al. 2016). In future work, it will be interesting to test the hypothesis that CCAR-1 affects germline apoptosis through the regulation of piRNAs and P granule function.

Investigation of genes involved in lifespan extension has been an area of interest in *C. elegans* research. In a previous study, we showed that knockdown of CCAR-1 affects an increase in the lifespan of wildtype N2 *C. elegans* at 23°C (Brunquell et al. 2018a). Here, we confirm this finding in wildtype N2 worms; however, with the use of a germline-defective worm strain, we see that CCAR-1 knockdown loses its lifespan extension capabilities at 25°C. Thus, the effect of CCAR-1 on lifespan regulation depends on the germline.

In summary, our experiments have established that CCAR-1 has a vital role in regulating the *C. elegans* germline. CCAR-1 regulates germline transcription, reproduction, DNA-damage induced apoptosis, and lifespan. We find it intriguing that CCAR-1 regulates various genes affiliated with P granules, membraneless organelles through which all germline mRNAs must pass through as they exit the nuclear pores. Are many of the germline-specific biological effects of CCAR-1 occurring through mRNA surveillance that occurs in the P granules? In future work, it will be important to unravel the precise mechanism of action of CCAR-1 in the germline.

## 7. ACKNOWLEDGEMENTS

The wildtype N2, CCAR-1Δ VC1029 (gk433), CCAR-1Δ MT7019 (n2667), CCAR-1Δ DM1153 (ra14), CCAR-1Δ DM1154 (ra5), PRG-1Δ WM161 (tm872), ACT-5∷YFP strain WS2170 (unc-119(ed-3)), GLP-1Δ CF1903 (e2144), strains were provided by the CGC, which is funded by NIH Office of Research Infrastructure Programs (P40 OD010440). This work was funded by NIH grant AG052149.

## REFERENCES

Bagijn, M.P., L.D. Goldstein, A. Sapetschnig, E.M. Weick, S. Bouasker et al., 2012 Function, targets, and evolution of Caenorhabditis elegans piRNAs. Science 337 (6094):574–578.

Basu, S., M. Barad, D. Yadav, A. Nandy, B. Mukherjee et al., 2020 DBC1, p300, HDAC3, and Siah1 coordinately regulate ELL stability and function for expression of its target genes. Proc Natl Acad Sci U S A 117 (12):6509–6520.

Berdichevsky, A., M. Viswanathan, H.R. Horvitz, and L. Guarente, 2006 C. elegans SIR-2.1 interacts with 14-3-3 proteins to activate DAF-16 and extend life span. Cell 125 (6):1165–1177.

Brunquell, J., R. Raynes, P. Bowers, S. Morris, A. Snyder et al., 2018a CCAR-1 is a negative regulator of the heat-shock response in Caenorhabditis elegans. Aging Cell 17 (5):e12813.

Brunquell, J., A. Snyder, F. Cheng, and S.D. Westerheide, 2017 HSF-1 is a regulator of miRNA expression in Caenorhabditis elegans. PLoS One 12 (8):e0183445.

Brunquell, J., J. Yuan, A. Erwin, S.D. Westerheide, and B. Xue, 2014a DBC1/CCAR2 and CCAR1 Are Largely Disordered Proteins that Have Evolved from One Common Ancestor. BioMed Research International 2014.

Close, P., P. East, A.B. Dirac-Svejstrup, H. Hartmann, M. Heron et al., 2012 DBIRD complex integrates alternative mRNA splicing with RNA polymerase II transcript elongation. Nature 484 (7394):386–389.

Ebbing, A., A. Vertesy, M.C. Betist, B. Spanjaard, J.P. Junker et al., 2018 Spatial Transcriptomics of C. elegans Males and Hermaphrodites Identifies Sex-Specific Differences in Gene Expression Patterns. Dev Cell 47 (6):801–813 e806.

Francois, S., C. D’Orlando, T. Fatone, T. Touvier, P. Pessina et al., 2012 Necdin enhances myoblasts survival by facilitating the degradation of the mediator of apoptosis CCAR1/CARP1. PLoS One 7 (8):e43335.

Fu, J., J. Jiang, J. Li, S. Wang, G. Shi et al., 2009 Deleted in breast cancer 1, a novel androgen receptor (AR) coactivator that promotes AR DNA-binding activity. J Biol Chem 284 (11):6832–6840.

Fu, R., Y. Zhu, X. Jiang, Y. Li, M. Zhu et al., 2018 CCAR-1 affects hemidesmosome biogenesis by regulating unc-52/perlecan alternative splicing in the C. elegans epidermis. J Cell Sci 131 (11).

Gartner, A., P.R. Boag, and T.K. Blackwell, 2008 Germline survival and apoptosis. WormBook:1–20.

Greer, E.L., T.J. Maures, A.G. Hauswirth, E.M. Green, D.S. Leeman et al., 2010 Members of the H3K4 trimethylation complex regulate lifespan in a germline-dependent manner in C. elegans. Nature 466 (7304):383–387.

Kanno, Y., S. Zhao, N. Yamashita, N. Saito, A. Ujiie et al., 2019 Cell Cycle and Apoptosis Regulator 1, CCAR1, Regulates Enhancer-Dependent Nuclear Receptor CAR Transactivation. Mol Pharmacol 95 (1):120–126.

Kim, J.E., J. Chen, and Z. Lou, 2008 DBC1 is a negative regulator of SIRT1. Nature 451 (7178):583–586.

Kinchen, J.M., J. Cabello, D. Klingele, K. Wong, R. Feichtinger et al., 2005 Two pathways converge at CED-10 to mediate actin rearrangement and corpse removal in C. elegans. Nature 434 (7029):93–99.

Lant, B., and W.B. Derry, 2013 Methods for detection and analysis of apoptosis signaling in the C. elegans germline. Methods 61 (2):174–182.

Lin, K., H. Hsin, N. Libina, and C. Kenyon, 2001 Regulation of the Caenorhabditis elegans longevity protein DAF-16 by insulin/IGF-1 and germline signaling. Nat Genet 28 (2):139–145.

Min, H., Y.H. Shim, and I. Kawasaki, 2016 Loss of PGL-1 and PGL-3, members of a family of constitutive germ-granule components, promotes germline apoptosis in C. elegans. J Cell Sci 129 (2):341–353.

Muthu, M., V.T. Cheriyan, and A.K. Rishi, 2015 CARP-1/CCAR1: a biphasic regulator of cancer cell growth and apoptosis. Oncotarget 6 (9):6499–6510.

Ou, C.Y., T.C. Chen, J.V. Lee, J.C. Wang, and M.R. Stallcup, 2014 Coregulator cell cycle and apoptosis regulator 1 (CCAR1) positively regulates adipocyte differentiation through the glucocorticoid signaling pathway. The Journal of biological chemistry 289 (24):17078–17086.

Povea-Cabello, S., M. Oropesa-Avila, P. de la Cruz-Ojeda, M. Villanueva-Paz, M. de la Mata et al., 2017 Dynamic Reorganization of the Cytoskeleton during Apoptosis: The Two Coffins Hypothesis. International journal of molecular sciences 18 (11).

Raynes, R., K.M. Pombier, K. Nguyen, J. Brunquell, J.E. Mendez et al., 2013 The SIRT1 modulators AROS and DBC1 regulate HSF1 activity and the heat shock response. PLoS One 8 (1):e54364.

Reinke, V., I.S. Gil, S. Ward, and K. Kazmer, 2004 Genome-wide germline-enriched and sex-biased expression profiles in Caenorhabditis elegans. Development 131 (2):311–323.

Seydoux, G., 2018 The P Granules of C. elegans: A Genetic Model for the Study of RNA-Protein Condensates. J Mol Biol 430 (23):4702–4710.

Wang, G., and V. Reinke, 2008 A C. elegans Piwi, PRG-1, regulates 21U-RNAs during spermatogenesis. Current biology : CB 18 (12):861–867.

Wang, X., Y. Zhao, K. Wong, P. Ehlers, Y. Kohara et al., 2009 Identification of genes expressed in the hermaphrodite germ line of C. elegans using SAGE. BMC Genomics 10:213.

Wang, Y., and H.A. Tissenbaum, 2006 Overlapping and distinct functions for a Caenorhabditis elegans SIR2 and DAF-16/FOXO. Mech Ageing Dev 127 (1):48–56.

Weick, E.M., and E.A. Miska, 2014 piRNAs: from biogenesis to function. Development 141 (18):3458–3471.

Wu, D.Y., C.Y. Ou, R. Chodankar, K.D. Siegmund, and M.R. Stallcup, 2014 Distinct, genome-wide, gene-specific selectivity patterns of four glucocorticoid receptor coregulators. Nuclear receptor signaling 12:e002.

Zhao, W., J.P. Kruse, Y. Tang, S.Y. Jung, J. Qin et al., 2008 Negative regulation of the deacetylase SIRT1 by DBC1. Nature 451 (7178):587–590.

